# Imaging invasive fungal growth and inflammation in NADPH oxidase-deficient zebrafish

**DOI:** 10.1101/703728

**Authors:** Taylor J. Schoen, Emily E. Rosowski, Benjamin P. Knox, David Bennin, Nancy P. Keller, Anna Huttenlocher

**Author notes:** Department of Biological Sciences, Clemson University, Clemson, South Carolina, USA. Corresponding author (AH). **Summary statement:** Live imaging reveals neutrophil NADPH oxidase activity limits fungal growth and excessive inflammation during *Aspergillus nidulans* infection in larval zebrafish.

## Abstract

Neutrophils are primary cells of the innate immune system that generate reactive oxygen species (ROS) and mediate host defense. Deficient phagocyte NADPH oxidase (PHOX) function leads to chronic granulomatous disease (CGD) that is characterized by invasive infections including those by the generally non-pathogenic fungus *Aspergillus nidulans*. The role of neutrophil ROS in this specific host-pathogen interaction remains unclear. Here, we exploit the optical transparency of zebrafish to image the effects of neutrophil ROS on invasive fungal growth and neutrophil behavior in response to *Aspergillus nidulans*. In a wild-type host, *A. nidulans* germinates rapidly and elicits a robust inflammatory response with efficient fungal clearance. PHOX-deficient larvae have increased susceptibility to invasive *A. nidulans* infection despite robust neutrophil infiltration. Expression of p22^phox^ specifically in neutrophils does not affect fungal germination but instead limits the area of fungal growth and excessive neutrophil inflammation and is sufficient to restore host survival in p22^phox^-deficient larvae. These findings suggest that neutrophil ROS limits invasive fungal growth and has immunomodulatory activities that contribute to the specific susceptibility of PHOX-deficient hosts to invasive *A. nidulans* infection.

## Introduction

Chronic granulomatous disease (CGD) is an inherited immunodeficiency caused by mutations in any of the five subunits comprising the phagocyte NADPH oxidase (PHOX) complex (p22^phox^, p40^phox^, p47^phox^, p67^phox^, p91^phox^) (Bedard and Krause, 2007) and is characterized by inflammatory disorders, granuloma formation, and recurrent or chronic bacterial and fungal infections such as invasive aspergillosis (IA) (King et al., 2016, Levine et al., 2005). While *Aspergillus* is typically innocuous to immunocompetent individuals, susceptibility to IA is increased in people with inherited immunodeficiency or medically-induced immunosuppression that impairs innate immune cell function, especially those with neutropenia (King et al., 2016, Segal et al., 2010). Relative to other inherited immune disorders, CGD is the most significant predisposing condition for developing IA (Blumental et al., 2011), and invasive aspergillosis is responsible for many of the infection-related mortalities of CGD patients (Henriet et al., 2012, Marciano et al., 2015), with *A. fumigatus* as the primary causative agent (Marciano et al., 2015). Despite its rare occurrence in other immunocompromised populations, *A. nidulans* infection is the second most frequent *Aspergillus* infection in CGD and is associated with higher morbidity and mortality rates than *A. fumigatus* (Henriet et al., 2012, Gresnigt et al., 2018). However, the underlying interactions between *A. nidulans* and the innate immune response that contribute to the unique susceptibility of CGD patients to *A. nidulans* infections remain unknown.

The pathological consequences of CGD are underpinned by the inability of CGD phagocytes to produce reactive oxygen species (ROS). Phagocytic ROS is reported to have both direct microbicidal and immunomodulatory functions, but which of these functions provide the dominant mechanism of host defense against *Aspergillus* remains unclear. *In vitro*, neutrophil-derived ROS can damage *Aspergillus* (Rex et al., 1990), but *Aspergillus* can counter oxidative stress through its own antioxidant pathways (Chang et al., 1998, Lambou et al., 2010, Wiemann et al., 2017). *In vivo*, mouse models of CGD are susceptible to *Aspergillus* infection, with both an impaired host defense and altered inflammatory response (Bonnett et al., 2006, Cornish et al., 2008, Morgenstern et al., 1997, Pollock et al., 1995). Furthermore, loss of PHOX activity causes aberrant inflammation in response to sterile injury, supporting a major role of PHOX in regulating inflammation independent of microbial control (Bignell et al., 2005, Morgenstern et al., 1997).

There is also a gap in understanding the cell-specific contribution of macrophage- and neutrophil-derived ROS in the innate immune response to *Aspergillus* infection. Host defense against inhaled *Aspergillus* conidia is mediated by the innate immune system (Balloy and Chignard, 2009) and in an immunocompetent host both macrophages and neutrophils can kill conidia or inhibit germination (Jhingran et al., 2012, Zarember et al., 2007), while neutrophils are primarily responsible for the destruction of hyphae post-germination (Diamond et al., 1978, Gazendam et al., 2016, Knox et al., 2014). There are varying reports on the importance of macrophage ROS for inhibiting conidial germination but there is consistent support for the role of macrophage ROS in regulating cytokines involved in neutrophil recruitment (Cornish et al., 2008, Grimm et al., 2013). In neutrophils, PHOX activity is dispensable for inhibiting germination *in vitro* (Zarember et al., 2007) but plays a minor role in germination inhibition and killing of *Aspergillus* spores *in vivo* (Cornish et al., 2008, Jhingran et al., 2012). Neutrophil ROS is also involved in regulating pro-inflammatory signals and neutrophils from CGD patients have increased expression of pro-inflammatory cytokines at a basal level as well as in response to pathogen challenge (Kobayashi et al., 2004, Smeekens et al., 2012). Together, the variable requirement of PHOX activity in restricting *Aspergillus* growth and the increased production of pro-inflammatory cytokines by CGD phagocytes support a role for PHOX activity in modulating neutrophil inflammation commonly associated with CGD. However, how neutrophil-specific ROS mediates fungal growth, fungal clearance, inflammation, and host survival in response to *A. nidulans* infection remains unclear.

In this study, we aimed to determine the neutrophil-specific role of PHOX in both clearing fungal burden and controlling inflammation and to identify characteristics of *A. nidulans* that allow it to cause disease specifically in CGD hosts by using an established *Aspergillus*-larval zebrafish infection model (Herbst et al., 2015, Knox et al., 2014, Rosowski et al., 2018). The transparency of zebrafish larvae and the availability of mutant and transgenic lines with fluorescently labeled phagocytes and cell-specific rescues allows us to observe both fungal burden and host inflammation over the course of a multi-day infection. Furthermore, zebrafish express functional PHOX in both neutrophils and macrophages (Brothers et al., 2011, Niethammer et al., 2009, Yang et al., 2012), making the larval zebrafish an attractive model for studying the cell specific roles of ROS in progression of infection.

We demonstrate that PHOX-deficient larvae (*p22^phox-/- (sa11798)^*) have increased susceptibility to *A. nidulans* infection, similar to human disease. Live imaging reveals that in a wild-type host, *A. nidulans* germinates faster, evokes a stronger immune response, and is cleared faster, compared to *A. fumigatus*. Global PHOX activity does not prevent conidial germination but reduces extensive invasive growth of *A. nidulans* hyphae and prevents excessive neutrophil recruitment to the infection site. Restoring PHOX activity in just neutrophils limits the area of invasive fungal growth and fully restores neutrophil recruitment and larval survival to wild-type levels. Our data demonstrate that *A. nidulans* elicits a distinct immune response that leads to both greater inflammation and hyphal-induced tissue damage in PHOX-deficient hosts and that neutrophil-derived ROS can limit both invasive fungal growth and inflammation.

## Results

### *p22P^phox-/- (sa11798)^* zebrafish larvae as a model of Chronic Granulomatous Disease (CGD)

To develop a zebrafish model of CGD, we began by characterizing embryos containing the *p22^phox^* allele sa11798 from the Sanger Zebrafish Mutation Project. This allele has a nonsense mutation that is predicted to cause the loss of 1 of 3 transmembrane domains of the p22 protein (Fig 1A) and will be referred to as *p22*^-/-^ herein. *p22*^-/-^ larvae had decreased p22 protein levels, as determined by Western blot analysis (Fig 1B). *p22*^-/-^ larvae also showed reduced survival in response to both *A. fumigatus* and *A. nidulans*, supporting its use as a model of CGD. Interestingly, *A. nidulans* caused more death than *A. fumigatus* in *p22*^-/-^ larvae, with a hazard ratio of 74.2 over infection in control larvae, compared to just 7.3 for *A. fumigatus*, demonstrating a specifically increased susceptibility to *A. nidulans* infection (Fig 1C). To determine if this *A. nidulans-induced* death was unique to a PHOX-deficient host, we also compared the ability of *A. nidulans* and *A. fumigatus* to cause disease in a generally immunosuppressed host. Wild-type larvae were infected with *A. nidulans* or *A. fumigatus* and treated with the glucocorticoid and general immunosuppressant dexamethasone or a DMSO vehicle control. In contrast to the *p22* mutant, *A. nidulans* caused less death in these dexamethasone-treated larvae than *A. fumigatus* (Fig 1D). Together, these data demonstrate the specific increase in susceptibility to *A. nidulans* in *p22* mutant larvae, providing a powerful tool to study the role of phagocyte ROS in specific host-pathogen interactions.

**Fig 1.**
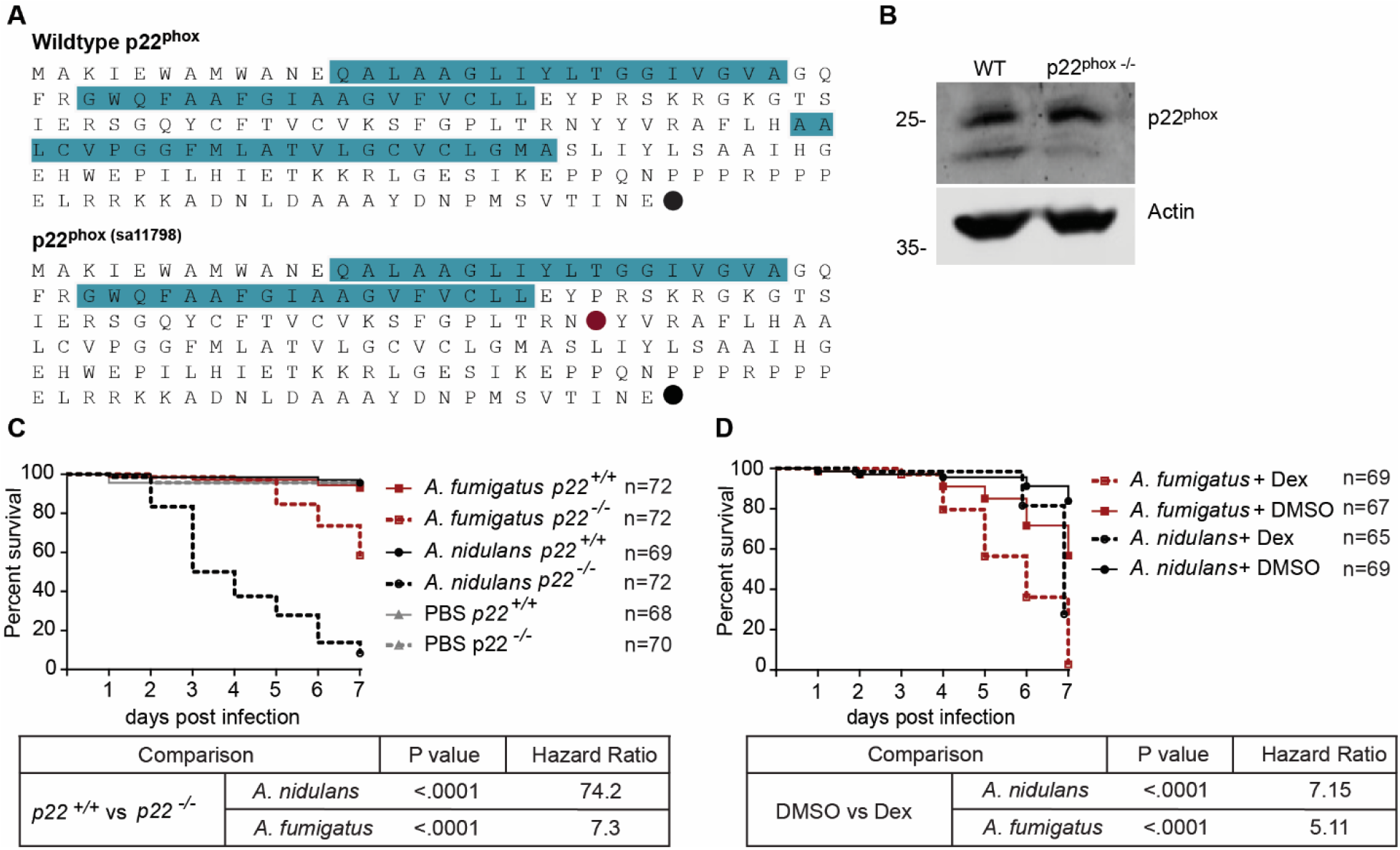
Characterization of p22^phox^ mutant zebrafish as a Chronic Granulomatous Disease model. (A) The *p22^phox^ sa11978* allele contains a point mutation that causes a premature stop codon (red circle: premature stop, black circle: normal stop) and putative loss of one of three transmembrane domains (each indicated by blue). (B) Western blot analysis using an antibody against human p22^phox^ with lysates from *p22*^+/+^ and *p22*^-/-^ larvae 2 days post fertilization. (C) *p22*^-/-^ and *p22*^+/+^ larvae were infected with *A. nidulans, A. fumigatus* or mock-infected with PBS. Average spore dose: *p22*^+/+^ with *A. nidulans* = 63, *p22*^-/-^ with *A. nidulans* = 62, *p22*^+/+^ with *A. fumigatus* = 59, *p22*^-/-^ with *A. fumigatus* = 64. (D) Wild-type larvae were infected with *A. nidulans* or *A. fumigatus* and treated with 10 μM dexamethasone or 0.1% DMSO vehicle control. Average spore dose: *A. nidulans* = 58, *A. fumigatus* = 75. Survival data (C and D) pooled from 3 independent replicates. P values and hazard ratios for survival experiments were calculated by Cox proportional hazard regression analysis.

### *A. nidulans* germinates faster and is cleared faster than *A. fumigatus* in wild-type hosts

Using live imaging, we first wanted to determine how fungal growth and host responses differ between *A. nidulans* and *A. fumigatus* infections in wild-type hosts. Germination of *Aspergillus* conidia to tissue-invasive hyphae is a critical driver of *Aspergillus* pathogenesis that triggers immune activation and fungal clearance by the host (Hohl et al., 2005, Rosowski et al., 2018). To determine if *A. nidulans* and *A. fumigatus* have different growth kinetics *in vivo*, we measured spore germination and hyphal growth in larval zebrafish. RFP-expressing *A. fumigatus* or *A. nidulans* spores were injected into the hindbrain of wild-type larvae and imaged by confocal microscopy at 6, 24, 48 and 72 hours post infection (hpi) to track *Aspergillus* germination and growth over the course of infection within individual larvae (Fig 2A-B). All *A. nidulans*-infected larvae contained germinated spores at 24 hpi but the percentage of larvae with persistent germinated spores decreased to 84% at 48 hpi and 47% at 72 hpi, suggesting clearance of *A. nidulans* by the host immune system (Fig 2C). In contrast, *A. fumigatus* germination was delayed until 48 hpi but the percentage of larvae with germination continued to increase up to 72 hpi with limited evidence of fungal clearance (Fig 2C). In addition to germination, we monitored invasive fungal growth as defined by the presence of lateral hyphal branching. The pattern of invasive growth followed a similar trend as germination. *A. nidulans*-infected larvae experienced faster development of invasive fungal growth and subsequent clearance relative to larvae infected with *A. fumigatus* (Fig 2D). While there was variability in fungal growth across individual larvae, the trend of early growth of *A. nidulans* followed by clearance was consistent, while all growth of *A. fumigatus* occurred later (Fig 2B).

**Fig 2.**
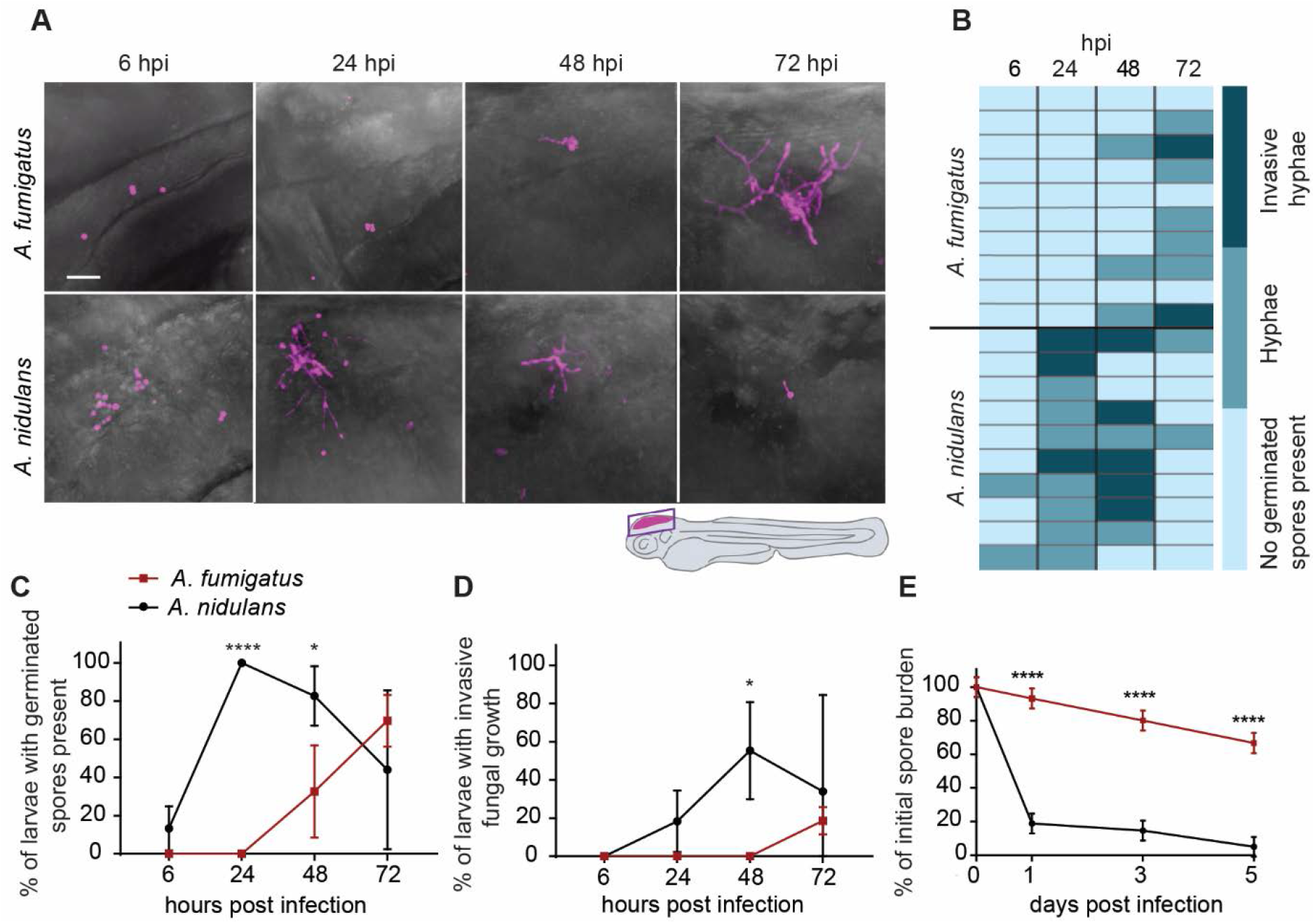
*A. nidulans* germinates faster and is cleared faster than *A. fumigatus* in a wild-type host. (A-D) Wildtype larvae were infected with RFP-labeled *A. nidulans* (*n*= 32) or RFP-labeled *A. fumigatus* (*n*= 32) and imaged by confocal microscopy. (A) Representative images of the same larvae followed throughout the 72 hour experiment. Images represent maximum intensity projections (MIP) of image z-stacks. Scale bar = 20 μm. (B) Heat map representing germination and hyphal growth status in infected larvae. Each row represents an individual larva, data are from one representative experiment. (C) Mean percentage of larvae containing germinated spores of *A. nidulans* or *A. fumigatus* throughout the 72 hour experiment, pooled from 3 independent replicates. (D) Mean percentage of larvae containing invasive fungal growth throughout the 72 hour experiment, pooled from 3 independent replicates. Invasive fungal growth was defined by the presence of hyphal branching. Data from C and D represent means ± SD, P values calculated by t-tests. (E) Individual larvae (*n* = 7-8 larvae per condition per day, 3 replicates) infected with *A. nidulans* or *A. fumigatus* were homogenized and plated to quantify fungal burden by CFU counts. Spore burden percentages are normalized to the mean CFU count from each condition on day 0. Spore burden data represents lsmeans ± SEM. P values calculated by ANOVA with Tukey’s multiple comparisons. Pooled from 3 independent replicates. Average spore dose: *A. nidulans* = 46, *A. fumigatus* = 42. * : P value < 0.05, **** : P value < 0.0001.

We further investigated differences in immune clearance of *A. fumigatus* and *A. nidulans* by quantifying the fungal burden in individual larvae through CFU plating experiments. Wild-type larvae were infected with *A. fumigatus* or *A. nidulans* spores and we measured CFUs at 0, 1, 3 and 5 dpi. A large percentage of the *A. fumigatus* spore burden persisted in the host up to 5 dpi, consistent with previous findings (Knox et al., 2014) while the *A. nidulans* spore burden dropped dramatically by 1 dpi (Fig 2E), providing further evidence that *A. nidulans* is cleared faster than *A. fumigatus* in wild-type larvae.

### *A. nidulans* infection activates a greater NF-κB response than *A. fumigatus*

Germination of *Aspergillus* reveals cell wall polysaccharides on the hyphal surface that contribute to immune activation (Henriet et al., 2016, Hohl et al., 2005). We hypothesized that differences in germination rate between *A. nidulans* and *A. fumigatus* would therefore result in differences in the inflammatory response following infection. To analyze NF-κB activation induced by *A. nidulans* and *A. fumigatus* infections *in vivo*, we utilized a NF-κB activation reporter line as a visual proxy of global pro-inflammatory activation (NF-κB RE:EGFP (Kanther et al., 2011)). At 1 dpi, *A. nidulans*-infected larvae had significantly more NF-κB reporter activity than *A. fumigatus*-infected larvae (Fig 3A-B), indicating that *A. nidulans* elicits a stronger host immune response early in infection as compared to *A. fumigatus*, likely contributing to its rapid clearance in a wild-type host.

**Fig 3.**
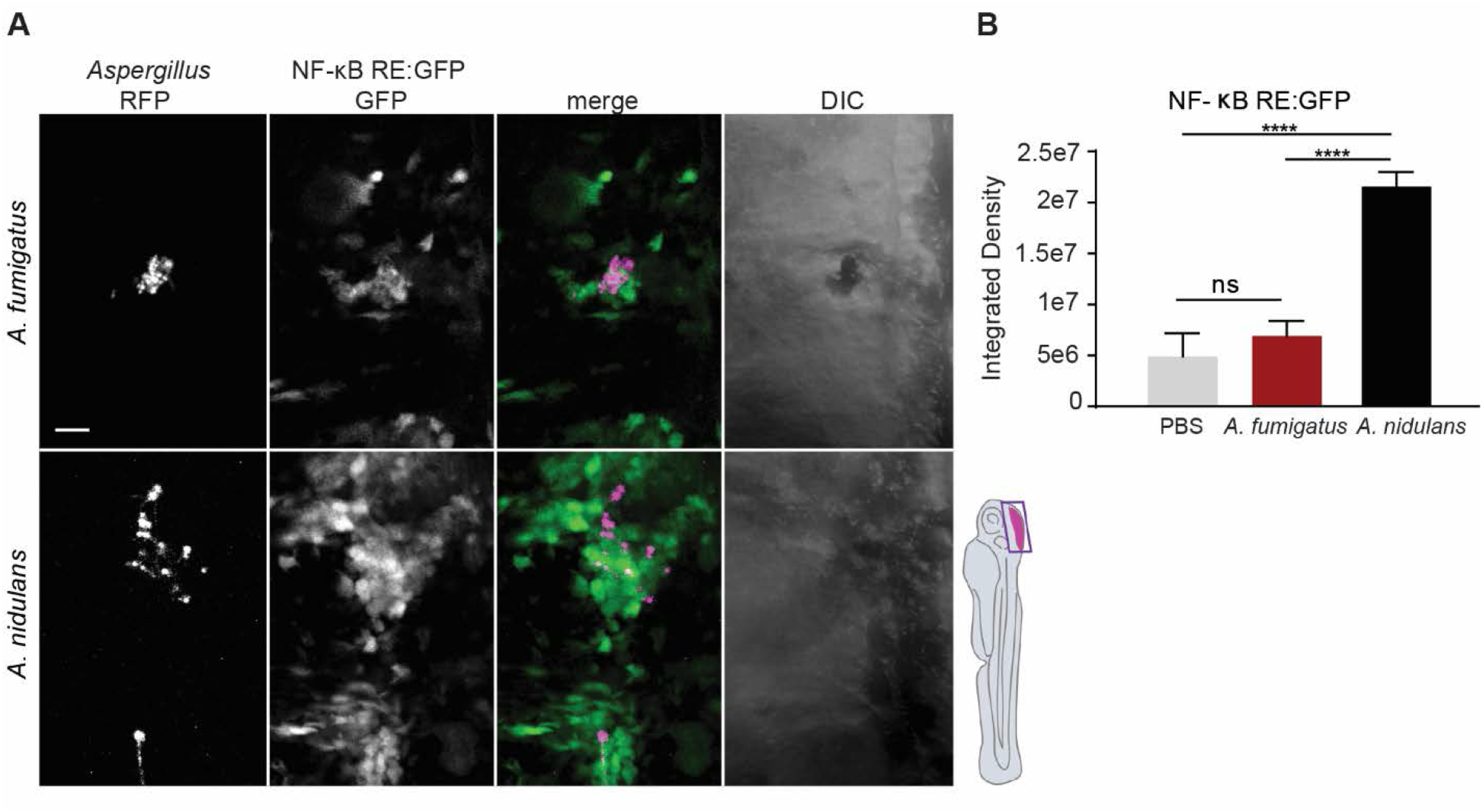
*A. nidulans* evokes a stronger NF-κB response than *A. fumigatus*. NF-κB RE:EGFP larvae were infected with RFP-expressing *A. nidulans* (*n*= 33), *A. fumigatus* (*n*= 32) or a PBS control (*n*= 13) and imaged by confocal microscopy at 1 dpi. (A) Images represent MIPs of image z-stacks. Scale bar = 20 μm. (B) NF-κB activation signal was quantified using the integrated density of a single z-slice from the hindbrain of individual larvae. Data represent lsmeans ± SEM from 3 pooled independent replicates. P values calculated by ANOVA with Tukey’s multiple comparisons. **** : P value < 0.0001.

### p22^phox^ controls *A. nidulans* invasive hyphal growth

We next determined whether the growth kinetics of *A. nidulans* are altered in *p22*^-/-^ larvae to begin to address why a PHOX-deficient host is more susceptible to *A. nidulans*. We infected *p22*^-/-^ and *p22*^+/-^ control larvae with RFP-expressing *A. nidulans* and imaged the larvae by confocal microscopy at 6, 24, 48, 72 and 96 hpi (Figs 4A-E, S3, Movies 1-2). In both *p22*^-/-^ and *p22*^+/-^ backgrounds, *A. nidulans* spore germination occurred by 24 hpi in 100% of larvae, with similar kinetics (Fig 4A-C). However, invasive hyphal growth was significantly increased, and fungal clearance was impaired by 48 hpi in *p22*^-/-^ larvae (Fig 4B-E). These data suggest a role for p22^phox^ in limiting hyphal growth post-germination rather than inhibiting the initiation of germination.

**Fig 4.**
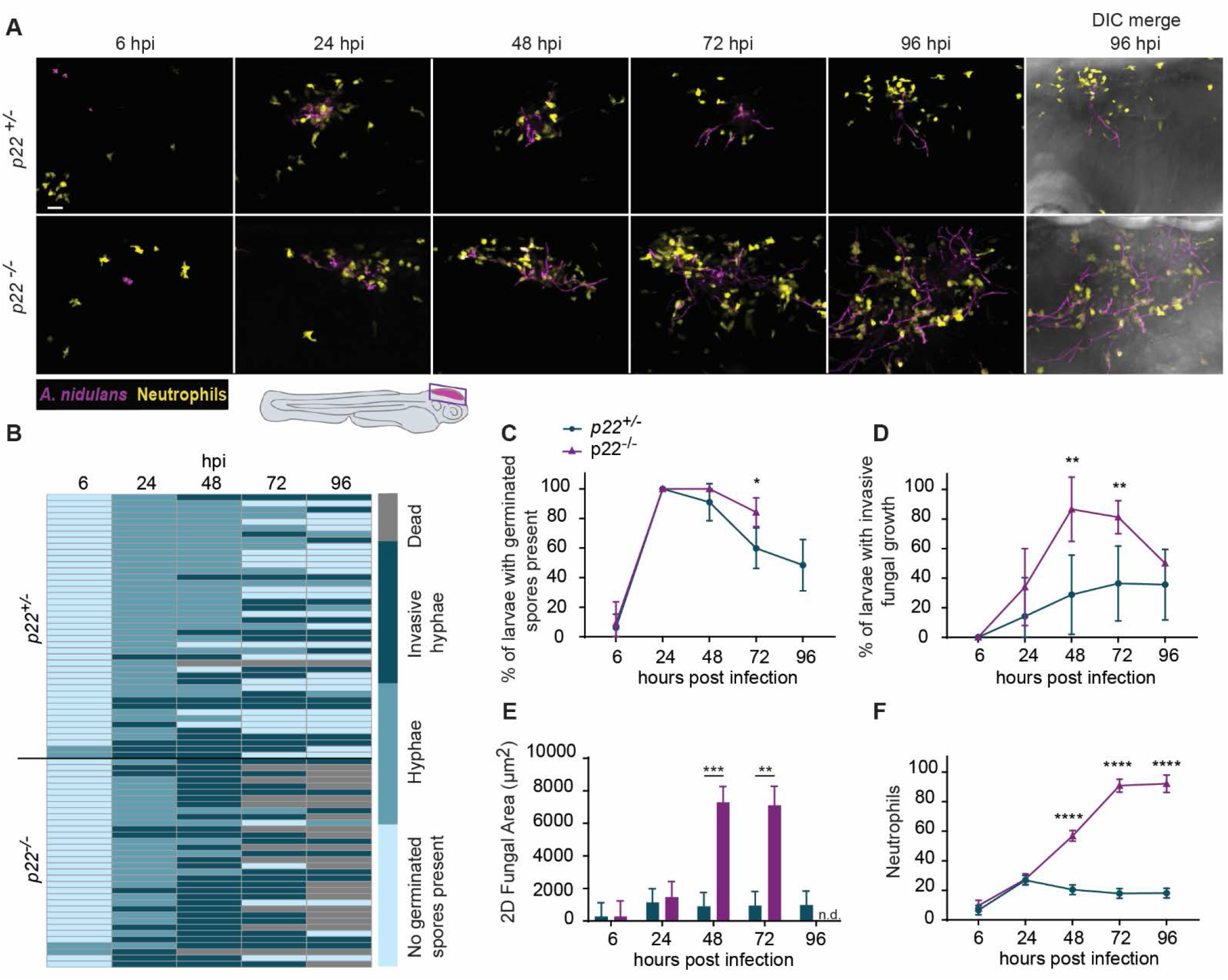
*p22*^-/-^ larvae infected with *A. nidulans* have higher fungal burdens and increased neutrophil recruitment. Neutrophil-labeled (*mpx:gfp*) *p22*^+/-^ and *p22*^-/-^ larvae were infected with RFP-labeled *A. nidulans* and imaged by confocal microscopy. (A) Representative images of the same larvae imaged throughout the 96 hour experiment. Images represent MIPs of image z-stacks. Scale bar = 20 μm. (B) Heat map representing germination and hyphal growth status in infected larvae. Data are compiled from 5 independent replicates. Each row represents an individual larva (*p22*^+/-^ *n* = 43, *p22*^-/-^ *n* = 34). (C) Mean percentage of larvae containing germinated spores, pooled from 4 independent replicates. (D) Mean percentage of larvae containing invasive fungal growth as determined by the presence of hyphal branching, pooled from 5 independent replicates. Data in C and D represent means ± SD and P values were calculated by t-tests. (E) Bars represent pooled lsmeans ± SEM of 2D fungal area from 5 independent replicates. At 96 hpi, only (13/34) *p22*^-/-^ larvae remained alive (see B), therefore data from that time point were excluded from C-E to better represent the group as a whole. (F) Quantification of neutrophil recruitment in the hindbrain of infected larvae. Data represent pooled lsmeans ± SEM of neutrophil counts from 5 independent replicates. P values calculated by ANOVA with Tukey’s multiple comparisons. * : P value < 0.05, ** : P value < 0.01, *** : P value < 0.001 **** : P value < 0.0001. n.d : no data.

### p22^phox^ controls neutrophil recruitment and resolution

In addition to fungal burden, PHOX can also modulate inflammatory cell recruitment. Because the primary innate immune cell recruited to hyphae are neutrophils (Diamond et al., 1978, Gazendam et al., 2016, Knox et al., 2014), we also imaged neutrophil infiltration over the course of *A. nidulans* infection in a labeled-neutrophil line (*mpx:gfp*) crossed with the *p22^phox^* mutant. *A. nidulans* recruited significantly more neutrophils in *p22*^-/-^ larvae as compared to *p22*^+/-^ larvae at 48, 72 and 96 hpi (Figs 4F, S1, Movies 1-2). While neutrophil recruitment in control hosts peaks at 24 hpi, recruitment in *p22*^-/-^ larvae continues to increase at least through 96 hpi and correlates with the presence of invasive hyphal growth.

### p22^phox^ negatively regulates inflammatory gene expression in response to *A. nidulans* infection

Previous studies have reported that PHOX-deficient hosts have increased expression of inflammatory cytokines (Henriet et al., 2012, Smeekens et al., 2012). To determine whether the increased neutrophil recruitment is associated with increased cytokine and chemokine expression, we performed RT-qPCR analyses of *il1b, il6, cxcl8-l1, cxcl8-l2*, and *tnfa* during *A. nidulans* infection in *p22*^+/+^ and *p22*^-/-^ larvae at 24 hpi. *p22*^-/-^ larvae exhibited variable levels of cytokine expression, but with statistically significant upregulation of *tnfa* and a trend towards increased expression of *il1b, il6, cxcl8-l2* in individual larvae compared to controls (Fig 5A-B). Taken together, our findings indicate that there is an increase in inflammatory mediators in *A. nidulans*-infected *p22*^-/-^ larvae early in infection as compared to *A. nidulans*-infected control larvae.

**Fig 5.**
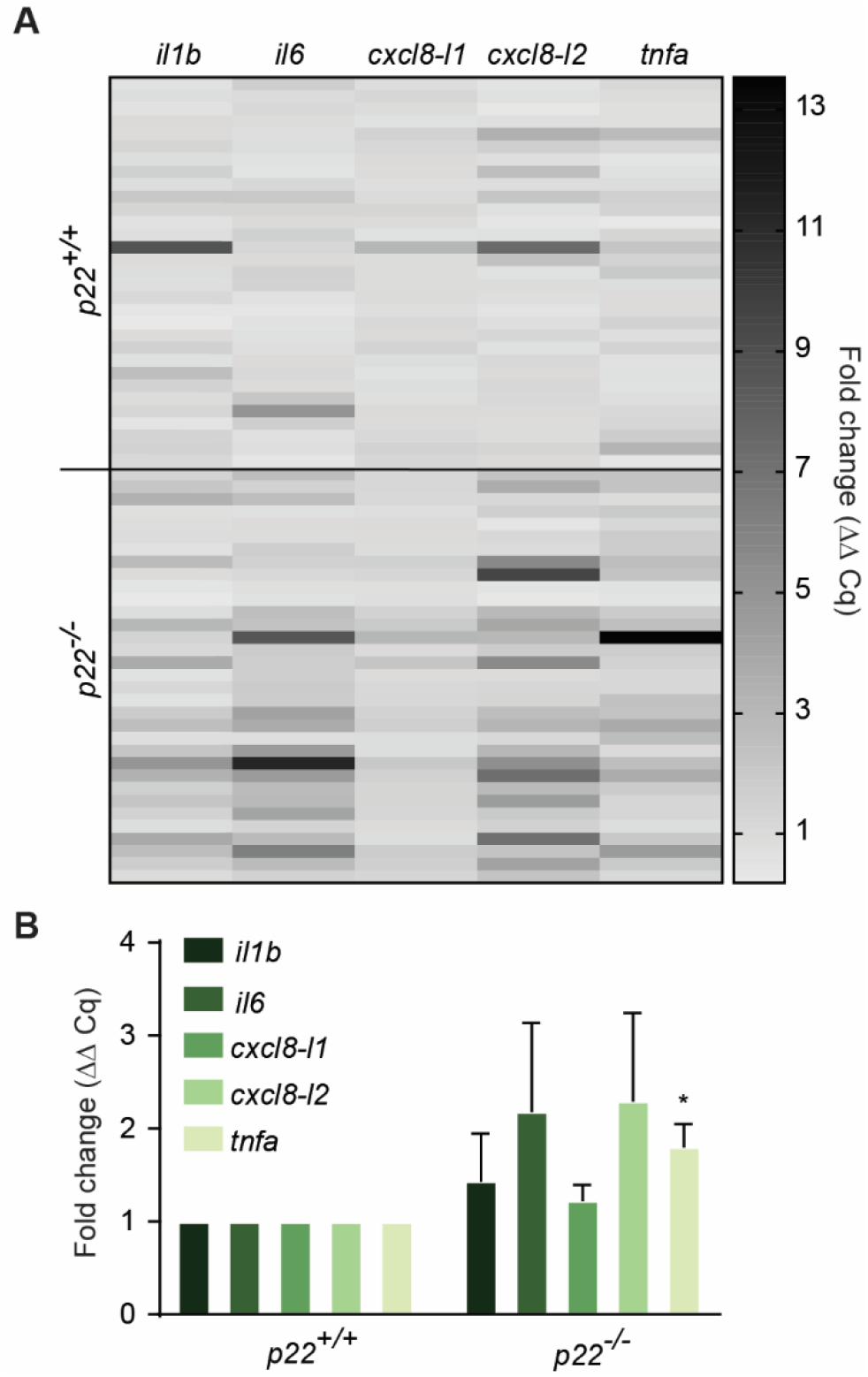
*A. nidulans* induces increased cytokine expression in *p22*^-/-^. RT-qPCR gene expression analysis of pro-inflammatory cytokines in *p22*^+/+^ and *p22*^-/-^ larvae infected with *A. nidulans* at 24 hpi. (A) Heat map representing the relative fold change of gene expression in individual larvae from 3 independent replicates as calculated by the ΔΔCq method. Each row represents an individual larva. (B) Bars represent the mean fold change in expression pooled from the individual larvae represented in (A) (*p22*^+/+^ *n* = 30, *p22*^-/-^ *n* =33). Average spore dose: *p22*^-/-^ = 77, *p22*^+/+^ = 75. P values for each cytokine were calculated by t-tests.* : P value < 0.05.

### Larvae with a neutrophil-specific deficiency in ROS production, but not neutropenic larvae, have increased susceptibility to *A. nidulans*

Our findings demonstrate increased neutrophil inflammation in response to *A. nidulans* in PHOX-deficient larvae, suggesting that neutrophils may play a key role in the pathogenesis of this infection. To further address the role of neutrophils and neutrophil ROS, we first utilized two different larval zebrafish models of human neutrophil deficiency, WHIM syndrome (*mpx:CXCR4-WHIM-GFP*) to model congenital neutropenia, and a model of leukocyte adhesion deficiency induced by a dominant negative Rac2D57N mutation in neutrophils (*mpx:rac2D57N*).While both models have impaired neutrophil migration to sites of infection (Deng et al., 2011, Walters et al., 2010), only the Rac2D57N model also has neutrophils that have an impaired oxidative burst (Ambruso et al., 2000, Williams et al., 2000). Larvae with WHIM and RacD57N mutations both showed impaired host survival in response to both *A. nidulans* and *A. fumigatus*. While there was no difference in survival of WHIM larvae infected with *A. fumigatus* or *A. nidulans* (Fig 6A), we observed decreased survival of Rac2D57N larvae infected with *A. nidulans* compared to those infected with *A. fumigatus* (Fig 6B), suggesting that neutrophil-derived ROS specifically mediates survival in response to *A. nidulans* infection.

**Fig 6.**
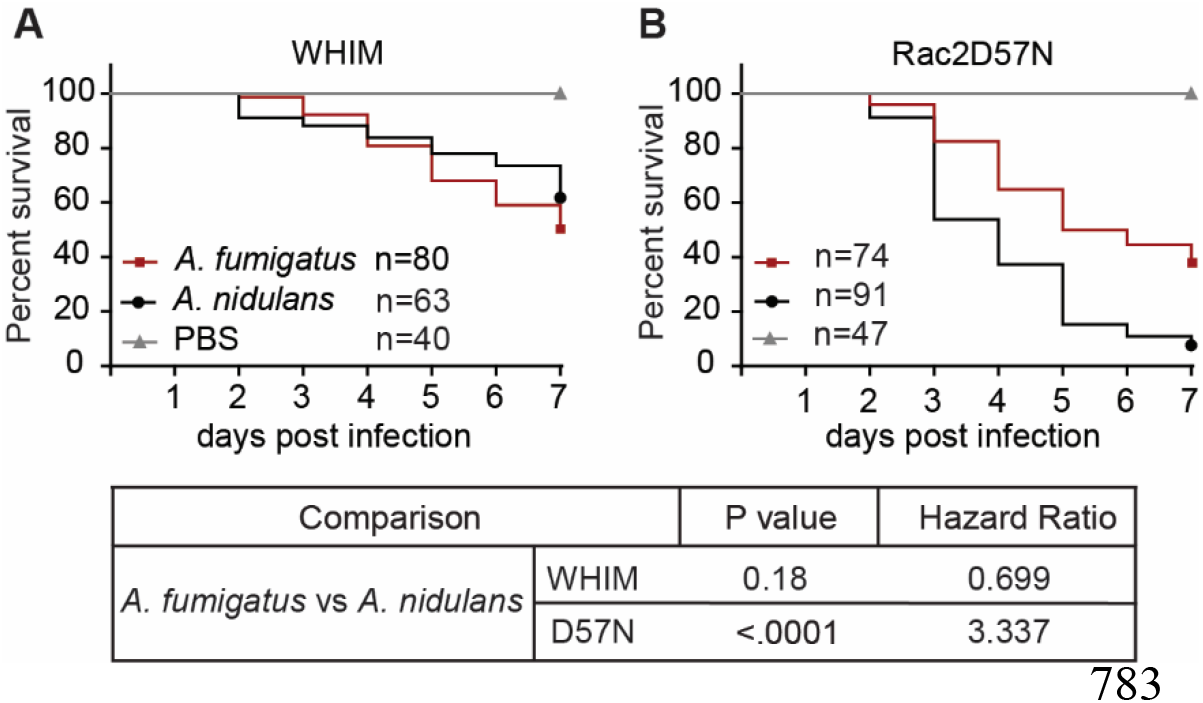
Rac2D57N larvae, but not WHIM larvae, have increased susceptibility to *A. nidulans*. Zebrafish models of the human neutrophil deficiencies WHIM syndrome (*mpx:CXCR4-WHIM-GFP*) and the dominant negative Rac2D57N mutation (*mpx:rac2D57N*) were infected with *A. nidulans* or *A. fumigatus*. (A) Survival analysis of WHIM larvae. Average spore dose: *A. nidulans* = 65, *A. fumigatus* = 54. (B) Survival analysis of Rac2D57N mutant larvae. Average spore dose: *A. nidulans* = 44, *A. fumigatus* =51. P values and hazard ratios calculated by Cox proportional hazard regression analysis.

### Neutrophil-specific expression of p22^phox^ rescues invasive fungal growth, excessive neutrophil inflammation and host survival in *p22*^-/-^ mutants

We therefore tested whether PHOX-derived ROS production in neutrophils alone was sufficient to mediate survival of *p22*^-/-^ larvae. We infected *p22*^-/-^; (*mpx:gfp*) and *p22*^-/-^ neutrophil rescue larvae that re-express functional p22^phox^ exclusively in neutrophils (*p22*^-/-^; *mpx:p22:gfp*) with RFP-expressing *A. nidulans* and imaged larvae by confocal microscopy at 6, 24, 48, 72, and 96 hpi (Figs 7A-F, 8A). *p22*^-/-^ larvae with p22^phox^ re-expressed just in neutrophils had a similar burden of germinated *A. nidulans* spores but have less extensive fungal growth than *p22*^-/-^ larvae, suggesting that neutrophil production of ROS limits the area of invasive fungal growth (Figs 7A-E, 8A). Furthermore, neutrophil-specific expression of functional p22^phox^ almost completely restored wild-type-like neutrophil recruitment to infection, suggesting that neutrophil-produced ROS may also limit excessive neutrophil inflammation (Figs 7F, S2). Because rescue of p22^phox^ in neutrophils alone improved both fungal burden and neutrophil inflammation in *p22*^-/-^ larvae, we next tested if neutrophil specific p22^phox^ expression improved overall host survival. Indeed, we found that neutrophil-specific expression of functional p22^phox^ rescued host survival back to wild-type levels (Fig 8B). These data demonstrate that neutrophil p22^phox^ contributes to clearance of hyphae and is sufficient for regulating excessive neutrophil recruitment. Control of both fungal growth and excessive inflammation allows for restoration of survival of larvae and suggests an important role of neutrophil-specific p22^phox^ in controlling *A. nidulans* infections.

**Fig 7.**
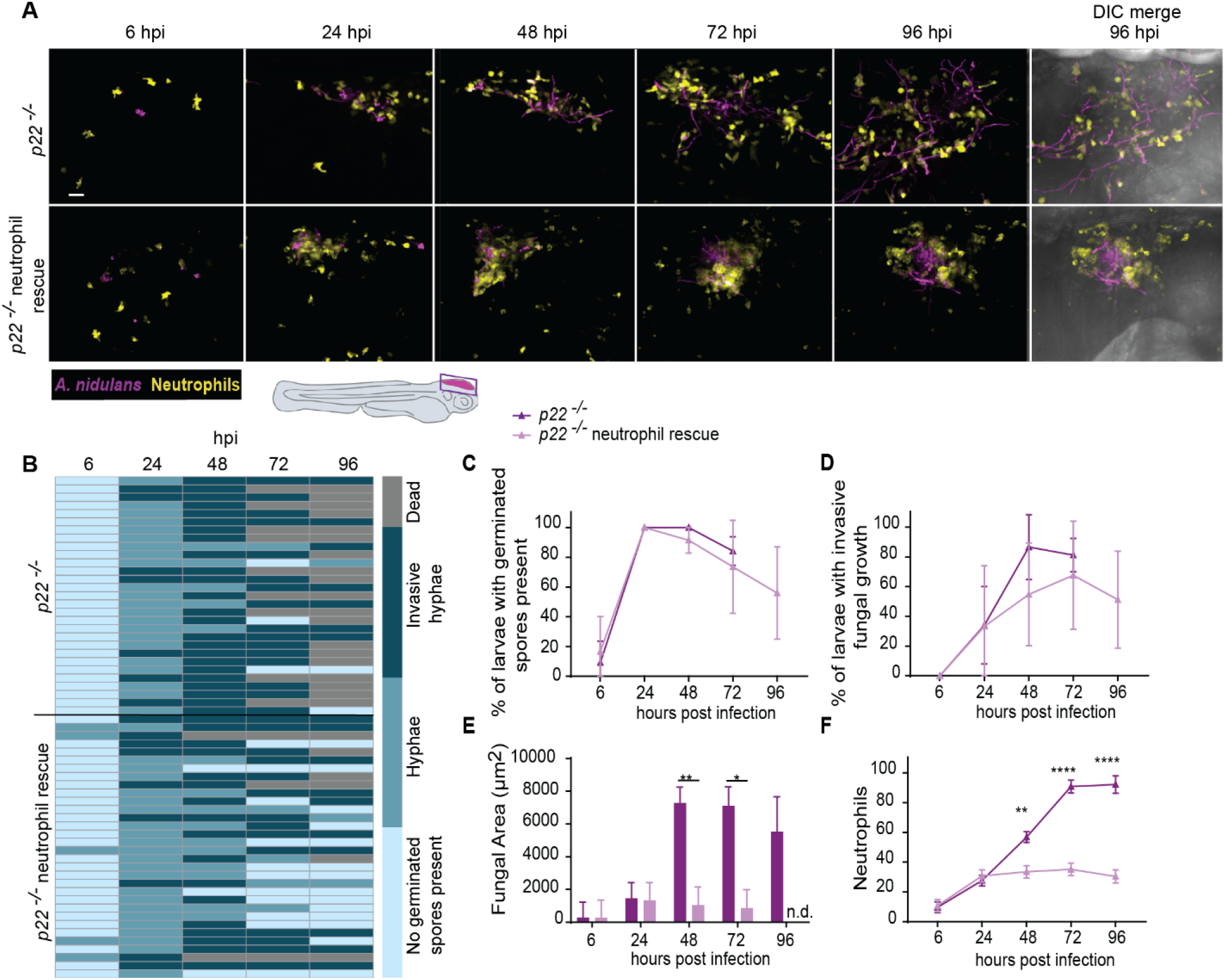
Neutrophil-specific expression of p22^phox^ rescues invasive fungal growth and excessive inflammation. *p22*^-/-^ larvae that re-express functional p22^phox^ in neutrophils (*p22*^-/-^; *mpx:p22:gfp*) and neutrophil-labeled *p22*^-/-^ larvae (*p22*^-/-^; *mpx:gfp*) were infected with RFP-labeled *A. nidulans* and imaged by confocal microscopy up to 96 hpi. *Note that analysis of p22 neutrophil-specific rescue larvae was performed in the same experiments described in Fig 4 and that the p22^-/-^ data in Fig 4 is repeated here for comparison with the neutrophil-specific rescue line*. (A) Images represent MIPs of image z-stacks. Scale bar = 20 μm. (B) Heat map representing germination and hyphal growth status in infected larvae from 5 independent replicates. Each row represents an individual larva (*p22*^-/-^ neutrophil rescue *n* = 27, *p22*^-/-^ *n* =34). (C) Mean percentage of larvae containing germinated spores, pooled from 5 independent replicates. (D) Mean percentage of larvae containing invasive fungal growth as determined by the presence of hyphal branching, pooled from 5 independent replicates. Data from C and D represent means ± SD, P values calculated by t-tests. (E) Bars represent pooled lsmeans ± SEM of 2D fungal area from 5 independent replicates. At 96 hpi, only (13/34) *p22*^-/-^ larvae remained alive (see B), therefore data from that time point were excluded from (C-E) to better represent the group as a whole. (F) Quantification of neutrophil recruitment in the hindbrain of infected larvae. Data represent pooled lsmeans ± SEM of neutrophil counts from 5 independent replicates. P values calculated by ANOVA with Tukey’s multiple comparisons.* : P value < 0.05, ** : P value < 0.01 **** : P value < 0.0001. n.d : no data.

**Fig 8.**
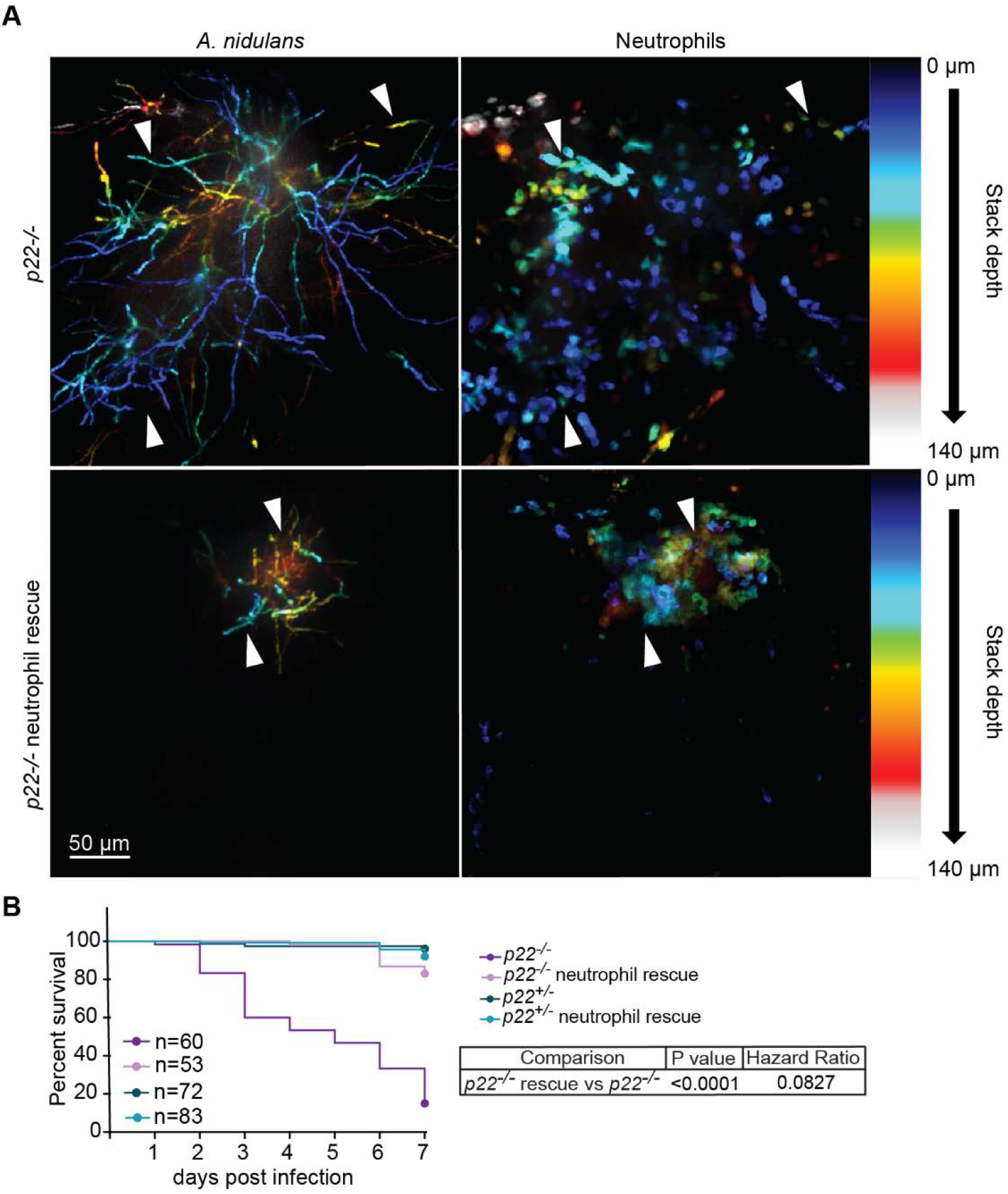
Neutrophil-specific expression of p22^phox^ rescues host survival. *p22*^-/-^ larvae that re-express functional p22^phox^ in neutrophils (*p22*^-/-^; *mpx:p22:gfp*) and neutrophil-labeled *p22*^-/-^ larvae (*p22*^-/-^; *mpx:gfp*) were infected with RFP-labeled *A. nidulans* and imaged by confocal microscopy up to 96 hpi. (A) Images represent depth-encoded MIPs of image z-stacks at 96 hpi. Color scale represents z-depth and location in the hindbrain in a 140 μm image z-stack. White arrows indicate examples of neutrophils occupying the same z-plane as hyphae. (B) Survival analysis of neutrophil labeled *p22*^-/-^ and *p22*^-/-^ neutrophil rescue larvae infected with *A. nidulans*. Average spore dose: *p22*^-/-^ = 77, *p22*^-/-^ neutrophil rescue= 83). P values and hazard ratios calculated by Cox proportional hazard regression analysis.

## Discussion

Here, we developed a PHOX-deficient zebrafish model to image the temporal and spatial dynamics of *A. nidulans* infection in a live, intact host. Few studies have addressed the unique susceptibility to the typically avirulent fungus *A. nidulans* in CGD. We find that *A. nidulans* is significantly more virulent than *A. fumigatus* in *p22*^-/-^ larvae but not in a generally immunocompromised or neutropenic host. Interestingly, *A. fumigatus* caused only a low level of death in *p22*^-/-^ larvae despite being the most frequently isolated fungus from CGD patients (King et al., 2016). The relatively weak virulence of *A. fumigatus* could be a result of strain variation, which can have a significant impact on host survival (Amarsaikhan et al., 2014, Knox et al., 2016, Kowalski et al., 2016, Rizzetto et al., 2013, Rosowski et al., 2018). Remarkably, rescuing p22^phox^ function specifically in neutrophils was sufficient to reduce invasive fungal growth but did not affect spore germination. It also fully restored neutrophil recruitment and host survival to wild-type levels, suggesting that death of *p22*^-/-^ larvae is a combined result of unconstrained fungal growth and damaging inflammation that is regulated by neutrophil production of ROS. To our knowledge, this is the first report of neutrophil ROS regulating host susceptibility to a specific fungal pathogen.

Using non-invasive live imaging techniques, we tracked germination and hyphal growth over the course of infection and saw that *A. nidulans* germinates faster and develops invasive hyphae faster than *A. fumigatus*, which both likely contribute to the clearance of *A. nidulans* in a wild-type host. Germination reveals immunogenic carbohydrates on the hyphal surface that induce phagocyte response to infection (Hohl et al., 2005), and we recently reported that faster germination and hyphal growth can increase fungal clearance by increasing immune activation and neutrophil-mediated killing (Rosowski et al., 2018). Similarly, we observe stronger NF-κB activation by *A. nidulans* than *A. fumigatus* at 24 hpi, concomitantly with greater germination and fungal growth, consistent with other reports that *A. nidulans* stimulates the immune system more than *A. fumigatus* (Gresnigt et al., 2018, Henriet et al., 2016, Smeekens et al., 2012). Considering this increased immune activation and the fact that CGD is defined by chronic inflammation, we hypothesize that in a wild-type host, this heightened immune response translates into rapid fungal clearance, but in a PHOX-deficient host it results in host-damaging inflammation.

Here, we observed both extensive hyphal growth and massive neutrophil recruitment to *A. nidulans*-infected p22^phox^-deficient larvae. The excessive neutrophil recruitment to *A. nidulans* infection observed in p22^phox^-deficient larvae is consistent with previous reports that PHOX activity regulates neutrophil inflammation during infection in mice and zebrafish (Mesureur et al., 2017, Segal et al., 2010) and could contribute to host death even in the absence of a functional PHOX complex, as neutrophils can damage host tissues by amplifying the response of other immune cells or by releasing enzymes and antimicrobial proteins that directly harm host tissue (Wang, 2018).

How does ROS regulate neutrophil recruitment and inflammation? We observed increased expression of pro-inflammatory cytokines in p22^phox^-deficient larvae in response to *A. nidulans*, recapitulating the enhanced cytokine production that is seen in human CGD patients (Kobayashi et al., 2004, Smeekens et al., 2012, Warris et al., 2003). We see consistent upregulation of *tnfa* and a trend towards increased *il6* and *cxcl8-l2* (IL-8) expression at 24 hpi, similar to observations in CGD mice (Morgenstern et al., 1997). TNFα and IL-6 mediate response to infection by promoting neutrophil recruitment (Hind et al., 2018) and ROS production (Figari et al., 1987). Additionally, CXCL8 is a strong neutrophil chemoattractant in humans (Hoffmann et al., 2002) and zebrafish (de Oliveira et al., 2013). Interestingly, zebrafish have 2 homologs of human CXCL8, L1 and L2. CXCL8-L1 and L2 can have distinct or redundant roles depending on the stimuli, including mediating neutrophil recruitment versus resolution of inflammation (de Oliveira et al., 2015, de Oliveira et al., 2013). Further study of the effects of these two isoforms in this CGD model may provide insight into whether the hyper-inflammation in this CGD model is due to enhanced neutrophil recruitment or impaired resolution.

Additionally, the specific susceptibility of Rac2D57N but not WHIM larvae to *A. nidulans* suggests that neutrophil ROS could be acting as a long-range signal to mediate inflammation, as neutrophils in WHIM larvae are not able to migrate to sites of infection (Walters et al., 2010) but can still produce ROS. Considering that ROS is diffusible through tissues (Niethammer et al., 2009) and the close association of macrophages and neutrophils during *Aspergillus* infection (Knox et al., 2017, Rosowski et al., 2018) it is possible that neutrophil-derived ROS could contribute to resolution of inflammation during infection by affecting macrophages. We previously reported that p22^phox^-deficient larvae have hyper-inflammation and defective neutrophil resolution at the site of sterile injury, partially due to impaired neutrophil reverse migration that is mediated by macrophages (Tauzin et al., 2014), demonstrating a signaling role for PHOX in regulating inflammation.

ROS has also been implicated as a direct microbial killer, but the ability of PHOX activity to kill *Aspergillus* has been debated. One argument against the importance of ROS in direct microbial killing in this specific CGD host-A. *nidulans* interaction is the finding that *A. nidulans* is not susceptible to killing by neutrophil-derived ROS *in vitro*, as compared to *A. fumigatus* (Henriet et al., 2011). In our experiments, we do observe significantly greater hyphal burden of *A. nidulans* in *p22*^-/-^ compared to *p22*^+/-^ larvae. However, the increased inflammation and resulting tissue damage in these larvae may also create a better environment for *A. nidulans* growth, making it difficult to parse apart these two roles for PHOX activity.

Our data support a larval zebrafish model with conserved p22^phox^ activity that is critical for defense against *A. nidulans* infection and contributes to host survival by mediating both fungal growth and damaging inflammation. Reconstitution of p22^phox^ in neutrophils supported the idea that there are cell-specific roles for PHOX activity during infection, however the full extent of phagocyte contributions to inflammation and fungal clearance warrants further investigation. For now, we think the full rescue of neutrophil recruitment and the partial rescue of invasive growth during *A. nidulans* infection contributes to host survival in 2 ways: 1) decreasing non-oxidative host tissue damage by neutrophils and 2) decreasing host tissue damage by invasive hyphae. Future work is needed to clarify if the neutrophilic inflammation observed in p22^phox^ deficient larvae damages host tissue and through what mechanisms. By exploiting the optical transparency and transgenic resources of the larval zebrafish model of CGD we and others are well-positioned to further investigate the mechanisms mediating PHOX-dependent regulation of inflammation and host survival.

## Methods

### Ethics statement

Animal care and use protocol M005405-A02 was approved by the University of Wisconsin-Madison College of Agricultural and Life Sciences (CALS) Animal Care and Use Committee. This protocol adheres to the federal Health Research Extension Act and the Public Health Service Policy on the Humane Care and Use of Laboratory Animals, overseen by the National Institutes of Health (NIH) Office of Laboratory Animal Welfare (OLAW).

### Fish lines and maintenance

Adult zebrafish and larvae were maintained as described previously (Knox et al., 2014). Larvae were anesthetized prior to experimental procedures in E3 water containing 0.2 mg/ml Tricaine (ethyl 3-aminobenzoate, Sigma). To prevent pigment formation in larvae during imaging experiments, larvae were maintained in E3 containing 0.2 mM N-phenylthiourea beginning 1 day post fertilization (dpf) (PTU, Sigma Aldrich). All zebrafish lines used in this study are listed in Table S1.

### Genotyping and line generation

Zebrafish containing the *p22^phox^* allele sa11798 were isolated through the Sanger Zebrafish Mutation Project, Wellcome Sanger Institute, and obtained from the Zebrafish International Resource Center (ZIRC). The sa11798 point mutation was detected using primers designed using the dCAPS method (Neff et al., 1998). Forward primer: p22_mutFor_MseI_F: 5’-CTTTTGGACCCCTGACCAGAAATTA-3’. Reverse primer: p22_mutFor_MseI_R: 5’-TGGCTAACATGAACCCTCCA-3’. The 211 bp PCR product was digested with MseI and analyzed on 3% agarose gel, with detected band sizes of: +/+ (115 bp, 96bp), +/- (115 bp, 96 bp, 74 bp) and -/- (115 bp, 74 bp). The neutrophil *p22^phox^* rescue line was genotyped with the same protocol as above, but the restriction pattern from these larvae contains an additional 84 bp product in all genotypes. For easier distinction of bands, the genotypes of these larvae were determined by analyzing digest products on 3% MetaPhor agarose gel. Neutrophil-specific restoration of p22^phox^ was achieved by crossing this *p22^phox^* sa11798 allele line with the *Tg*(*mpx:cyba:gfp*) line (Tauzin et al., 2014). The *Tg(mpx:cyba:gfp); p22^phox-/-^* is also referred to as *mpx:p22:gfp* in the results for simplicity. Larvae expressing GFP in neutrophils were selected to grow up and establish a stable line.

### *Aspergillus* strains and growth conditions

All *Aspergillus* strains used in this study are listed in Table S2. All strains were grown on solid glucose minimal medium (GMM). *A. fumigatus* was grown at 37°C in darkness and *A. nidulans* was grown at 37°C in constant light to promote asexual conidiation. Conidial suspensions for microinjection were prepared using a modified protocol from Knox et al., 2014 that includes an additional filtration step to eliminate hyphal fragments and conidiophores of *A. nidulans* from the suspension. For consistency, the additional filtration step was also used for isolation of *A. fumigatus* conidia. After *Aspergillus* was grown for 3-4 days on solid GMM after being plated at a concentration of 1 x 10^6^ conidia/10 cm plate, fresh conidia were harvested in 0.01% Tween water by scraping with an L-spreader. The spore suspension was then passed through sterile Miracloth into a 50 mL conical tube and the volume was adjusted to 50 mL with 0.01% Tween. The spore suspension was centrifuged at 900 x g for 10 minutes at room temperature and the spore pellet was re-suspended in 50 mL 1x PBS. The spore suspension was then vacuum filtrated using a Buchner filter funnel with a glass disc containing 10-15 μm diameter pores. The filtered suspension was centrifuged at 900 x g for 10 minutes and re-suspended in 1 mL 1x PBS. Conidia were counted using a hemacytometer and the concentration was adjusted to 1.5 x 10^8^ spores/mL. Conidial stocks were stored at 4°C and used up to 1 month after harvesting.

### Western blotting

For western blotting, 50-100 2 dpf larvae were pooled and de-yolked in calcium-free Ringer’s solution with gentle disruption with a p200 pipette. Larvae were washed twice with PBS and stored at −80°C until samples were lysed by sonication in 20mM Tris pH 7.6, 0.1% Triton-X-100, 0.2 mM Phenylmethylsulfonyl fluoride (PMSF), 1 μg/mL Pepstatin, 2 μg/mL Aprotinin, and 1 μg/mL Leupeptin at 300 μL per 100 larvae while on ice and clarified by centrifugation. Protein concentrations were determined using a bicinchoninic acid protein assay kit (Thermo Fisher Scientific), according to the manufacturer’s instructions. Approximately 100 μg total protein were loaded on 6-20% gradient SDS-polyacrylamide gels and transferred to nitrocellulose. Zebrafish p22^phox^ was detected using an antibody against full-length human p22^phox^ (p22-phox (FL-195): sc-20781, Santa Cruz Biotechnology, 1:500 dilution). Western blots were imaged with an Odyssey Infrared Imaging System (LI-COR Biosciences).

### Spore microinjections

Anesthetized 2 dpf larvae were microinjected with conidia into the hindbrain ventricle via the otic vesicle as described in Knox et al., 2014. 1% phenol red was mixed in a 2:1 ratio with the conidial suspension so the inoculum was visible in the hindbrain after injection. After infection larvae were rinsed 3x with E3 without methylene blue (E3-MB) to remove the Tricaine solution and were then transferred to individual wells of a 96-well plate for survival experiments. For imaging experiments larvae were kept in 35 mm dishes or 48-well plates. Survival was checked daily for 7 days and larvae with a heartbeat were considered to be alive. We aimed for an average spore dose of 60 spores and the actual spore dose for each experiment was monitored by CFU counts and reported in the figure legends.

### CFU counts

To quantify the initial spore dose and fungal burden in survival and fungal clearance assays, anesthetized larvae were collected immediately after spore injection and placed in individual 1.5 mL micro-centrifuge tubes in 90 μl of 1x PBS with 500 μg/ml Kanamycin and 500 μg/ml Gentamycin. Larvae were homogenized using a mini-bead beater for 15-20 seconds and the entire volume of the tube was plated on GMM solid medium. Plates were incubated at 37°C for 2-3 days and CFUs were counted. At least 7-8 larvae were used for each condition, time point and replicate. For quantifying the percent of initial spore burden, the number of CFUs were normalized to the average CFU count on day 0.

### Drug treatments

We used dexamethasone (Sigma) to induce general immunosuppression of wildtype larvae. Dexamethasone was re-constituted to 10 mM with DMSO for a stock solution and stored at −20°C. Immediately following spore microinjection, larvae were treated with 10 μM dexamethasone or DMSO (0.01%) and the larvae remained in the drug bath for the entirety of the survival experiment.

### RT-qPCR

RNA was extracted from individual larvae with 100 μl TRIzol reagent (Invitrogen) and cDNA was synthesized with SuperScript III RT and oligo-dT (Invitrogen). cDNA was used as the template for qPCR using FastStart Essential Green DNA Master (Roche) and a LightCycler96 (Roche). Data was normalized to *rps11* using the ΔΔCq method (Livak and Schmittgen, 2001). The mean of ΔΔCq values across multiple individual larvae was used to calculate the fold changes displayed in Figure 5B. Fold changes therefore represent pooled data collected from 33 individual *p22*^-/-^ and 30 *p22*^+/+^ larvae over three experiments. All qPCR primers are listed in Table S3.

### Live imaging

Pre-screening of larvae with fluorescent markers was performed on a zoomscope (EMS3/SyCoP3; Zeiss; Plan-NeoFluar Z objective). Multi-day imaging experiments were performed on a spinning disk confocal microscope (CSU-X; Yokogawa) with a confocal scanhead on a Zeiss Observer Z.1 inverted microscope, Plan-Apochromat NA 0.8/20x objective, and a Photometrics Evolve EMCCD camera. Images were acquired using ZEN software (Zeiss). Larvae imaged at multiple time points were kept in 48-well plates in E3-MB with PTU. For imaging, larvae were removed from the 48-well plate and anesthetized in E3-MB with Tricaine before being loaded into zWEDGI chambers (Huemer et al., 2017). Larvae were positioned so that the hindbrain was entirely visible during imaging. All z-series images were acquired in 5 μm slices. After imaging, the larvae were rinsed with E3-MB to remove Tricaine and were returned to the 48-well plate with E3-MB with PTU. To image GFP signal in the *Tg*(*NF-κB RE:GFP*) line, larvae were anesthetized and positioned at the bottom of a glass-bottom dish in 1% low-melting point agarose. Images were acquired on a laser-scanning confocal microscope (FluoView FV1000; Olympus) with an NA 0.75/20x objective and FV10-ASW software (Olympus).

### Image analysis and processing

All displayed images in Figs 1-7 represent maximum intensity projections of z-series images that were generated using Imaris software. The brightness and contrast were adjusted for displayed images also using Imaris. For analysis of germination and invasive growth, all images were viewed as z-stacks and maximum intensity projections in Zen software (Zeiss) to score the presence of germination and hyphae. For measuring GFP signal in *Tg*(*NF-κB RE:GFP*) larvae, images were analyzed in Fiji. A single z-slice from the middle of the hindbrain was used to measure GFP signal intensity. The measurement area and integrated density of GFP signal measured inside an ROI encompassing the hindbrain, with the ROI determined manually from the brightfield image. No alterations were made to the images prior to analysis. 2D fungal area was measured by manually thresholding the RFP signal from *A. nidulans* and measuring the area of RFP signal. Fungal area measurements were made from maximum intensity projections of z-stacks in Fiji. Neutrophil recruitment during infection was analyzed by manually counting neutrophils within the hindbrain and neutrophils in contact with hyphae in cases where hyphae grew outside of the hindbrain. In addition to counting cells, neutrophil recruitment was also analyzed by manually thresholding the GFP signal from neutrophils and measuring the area of GFP signal within the hindbrain and neutrophils in contact with hyphae in cases where hyphae grew outside of the hindbrain. Both counting and GFP-signal area measurements were made from maximum intensity projections of z-stacks in Fiji. Movies 1 and 2 were prepared and brightness and contrast of each channel were adjusted in Fiji. Depth-encoded maximum intensity projections of z-stacks were generated in Fiji.

### Statistical analyses

All experiments and statistical analyses represent at least 3 independent replicates. The number of replicates for each experiment is indicated in the figure legends. Survival experiments were analyzed using Cox proportional hazard regression analysis with the experimental condition included as a group variable, as previously described (Knox et al., 2014). The pair-wise P values, hazard ratios, and experimental Ns are displayed in each figure. For analysis of spore germination and invasive growth pair-wise comparisons were made using student’s T-tests and data represent means ± standard deviation. Integrated densities of GFP signal, RFP signal, neutrophil counts, and spore burden represent least-squared adjusted means ± standard error of the mean (lsmeans±SEM) and were compared by ANOVA with Tukey’s multiple comparisons. Fold changes in gene expression in *p22*^-/-^ larvae were compared to the normalized control value of 1 using one-sample t-tests. Cox proportional hazard regression, least-squared adjusted means and ANOVA analyses were performed using R version 3.4.4. T-tests and graphical representations were done in GraphPad Prism version 7.

## Acknowledgements

We thank members of the Huttenlocher and Keller labs for helpful discussions of the research and manuscript. We thank Jens Eickhoff for guidance on and assistance with statistical analyses.

## Data availability

All relevant data are within the paper and its Supporting Information files.

